# The HRA Organ Gallery Affords Immersive Superpowers for Building and Exploring the Human Reference Atlas with Virtual Reality

**DOI:** 10.1101/2023.02.13.528002

**Authors:** Andreas Bueckle, Catherine Qing, Shefali Luley, Yash Kumar, Naval Pandey, Katy Börner

## Abstract

The Human Reference Atlas (HRA, https://humanatlas.io) funded by the NIH Human Biomolecular Atlas Program (HuBMAP, https://commonfund.nih.gov/hubmap) and other projects engages 17 international consortia to create a spatial reference of the healthy adult human body at single-cell resolution. The specimen, biological structure, and spatial data that define the HRA are disparate in nature and benefit from a visually explicit method of data integration. Virtual reality (VR) offers unique means to enable users to explore complex data structures in a threedimensional (3D) immersive environment. On a 2D desktop application, the 3D spatiality and real-world size of the 3D reference organs of the atlas is hard to understand. If viewed in VR, the spatiality of the organs and tissue blocks mapped to the HRA can be explored in their true size and in a way that goes beyond traditional 2D user interfaces. Added 2D and 3D visualizations can then provide data-rich context. In this paper, we present the HRA Organ Gallery, a VR application to explore the atlas in an integrated VR environment. Presently, the HRA Organ Gallery features 55 3D reference organs,1,203 mapped tissue blocks from 292 demographically diverse donors and 15 providers that link to 5,000+ datasets; it also features prototype visualizations of cell type distributions and 3D protein structures. We outline our plans to support two biological use cases: **on-ramping novice and expert users to HuBMAP data available via the Data Portal** (https://portal.hubmapconsortium.org), and **quality assurance/quality control (QA/QC) for HRA data providers**. Code and onboarding materials are available at https://github.com/cns-iu/ccf-organ-vr-gallery#readme.

## Introduction

We present the HRA Organ Gallery, a VR application implemented to explore the 3D organ models of the HRA in their true scale, location, and spatial relation to each other. We focus on two use cases: **on-ramping novice and expert users to the HuBMAP data available via the HuBMAP Data Portal** and **quality assurance/quality control (QA/QC) for HRA data providers**. **Figure 1** shows how users typically see the HRA via the Exploration User Interface(EUI) (Börner et al., 2022), vs. how they see it within the HRA Organ Gallery using a Meta Quest Pro.

**Figure 1.**
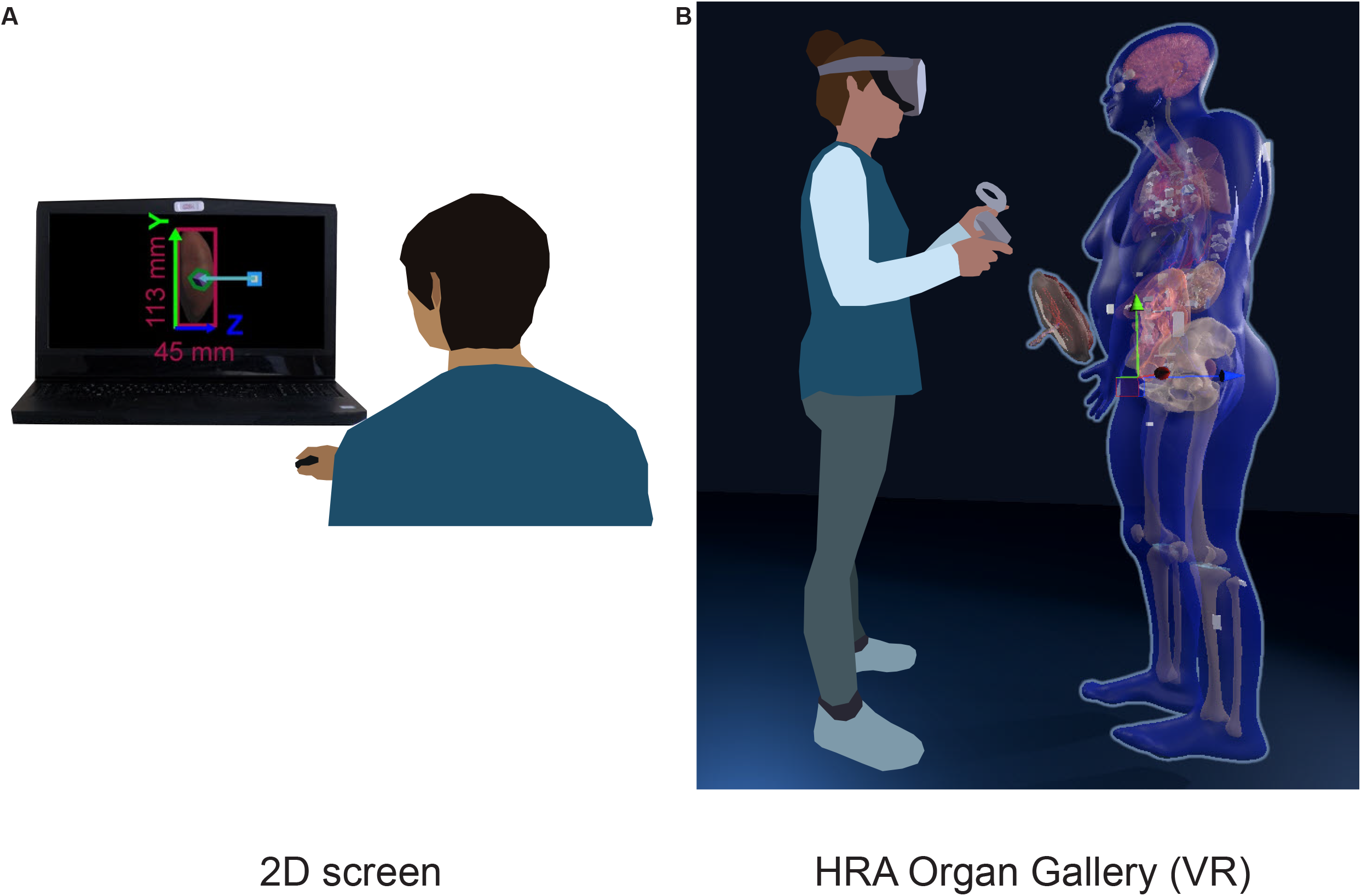
2D vs. VR. **(A)**: A user looks at the HRA with the Exploration User Interface (https://portal.hubmapconsortium.org/ccf-eui) in a standard-size browser window on a 17-in display. **(B)**: The HRA Organ Gallery allows the user to view the organs, tissue blocks, and cell type counts of the HRA in true scale using immersive technology (VR).

### Human Reference Atlas data

The HRA effort (Börner et al., 2021) led by HuBMAP (HuBMAP Consortium et al., 2019) is a close collaboration across 17 international consortia to map the adult healthy human body at single-cell resolution. The HRA consists of three major data types: **biological structure data**, detailing the relationship between anatomical structures (AS) inside human organs, the cell types (CT) located in them, and the biomarkers (B) that characterize them; (2) **spatial data** that defines entities inside a 3D reference system; and (3) **specimen data** describing donors in terms of their demographic and health markers, such as age, sex, ethnicity, and body mass index (BMI). **Biological structure data** is captured via Anatomical Structures, Cell Types, and Biomarker (ASCT+B) tables, which are compiled by experts to detail the cell types and biomarkers typically found in human anatomy (Börner et al., 2021). The recent HRA v1.3 release from December 2022 features 2,703 anatomical structures, 703 cell types, and 1,502 biomarkers. In synchrony with the ASCT+B table effort, there exists a matching set of 55 3D reference organs (Kristen Browne et al., 2022), mostly modeled after the Visible Human Project data provided by the National Library of Medicine (Spitzer et al., 1996), that comprise the **spatial data** of the HRA. The anatomical structures in the ASCT+B tables are linked to anatomical structures in the 3D reference organs via a crosswalk (Bruce W. Herr II, 2022). Using the Registration User Interfaces (RUI) and the EUI (Bueckle et al., 2021, 2022; Börner et al., 2022) 3D reference organs are used to register 3D tissue blocks from diverse donors into the HRA. The **specimen data** for each tissue block is tracked to support filter, search, and exploration by donor demographics using the CCF Ontology (Herr et al., 2022).

The **disparate data** defining the HRA benefits from a visually explicit method of data integration. VR offers unique means to explore spatial and abstract data in a unified, immersive, and presence-enhancing environment beyond traditional WIMP interfaces, i.e., windows, icons, menu, pointer (Van Dam, 1997). While exploring 3D reference organs and tissue blocks on a 2D screen can be learned (Bueckle et al., 2021, 2022), many users have issues interacting with 3D objects on a 2D screen.

### Related work

Recent advancements in VR development have led to an interest in applying VR to biomedical research and practice. VR allows dynamic exploration and enables viewers to enter visualizations at various viewpoints (Camp et al., 1998), and to create visualizations of intricate molecular structures and biomolecular systems (Chavent et al., 2011; Gill and West, 2014; Trellet et al., 2018; Wiebrands et al., 2018). These virtual displays offer seamless and dynamic user interactions with the 3D models.

In recent years, researchers have also explored VR as a method for users to engage with virtual models of human anatomy, and thus a tool for medical education (Marks et al., 2017; Moro et al., 2017; Ammanuel et al., 2019; Maresky et al., 2019; Liimatainen et al., 2021). VR has additionally been lauded as a valuable spatial instruction tool, as demonstrated by Nicholson et al.’s interactive ear model that users can zoom in and out of and smoothly rotate (Nicholson et al., 2006). Most previous research, however, has only investigated the implementation of one to a few 3D models; no work has developed and displayed 3D organ models in a virtual environment at the scale of the present work.

Not only does VR present advantages for 3D modeling, it also offers novel means of exploring and presenting data. For instance, information-rich virtual environments (IRVEs)(Bowman et al., 2003) are effective presentations of combined perceptual and abstract information (Bowman et al., 1998, 1999). Moreover, IRVEs are utilized increasingly for data visualization, leading to the establishment of Immersive Analytics (IA) in recent years as a field studying the use of VR for data analysis, visual analytics, and information visualization (Chandler et al., 2015; Cordeil et al., 2017b; Billinghurst et al., 2018; Batch et al., 2020). A distinct advantage of VR for visualizing scientific data alongside abstract data is its ability to preserve and present spatial dimensions of data in an immersive way. IRVEs also offer a vast design space, enabling users to be placed ‘inside’ the data. Embodiment is another captivating feature of IVEs, as users can directly manipulate, explore, and rearrange virtual objects mapped to data artifacts (Dourish, 2001). This concept is exemplified in works such as ImAxes (Cordeil et al., 2017a), which allows users to manipulate virtual axes to create data visualizations. Because VR supports spatial reasoning, intuitive interaction, and spatial memory, it is frequently used to support data exploration (Marriott et al., 2018). Resultantly, VR makes complex data more accessible and understandable (Lin et al., 2013).

VR offers a unique method of visualizing and interacting with the diverse data types that constitute the HRA in a unified 3D environment. The design of the HRA Organ Gallery uniquely focuses on enriching the analysis experience with presence and immersion while simultaneously preserving the spatiality of biological data. Not only is the HRA Organ Gallery the first open-source immersive application to allow users to view, explore, and analyze 55 3D reference organs in an integrated VR environment, it is the first such application to also integrate varying types of data and data visualizations while affording users with a broad range of actions (e.g., select cell types in anatomical structures).

## Method

### Use cases

HRA Organ Gallery research and development are driven by two concrete bioinformatic use cases:

#### (1) On-ramping of novice and expert users

The HuBMAP Data Portal (Human BioMolecular Atlas Program, 2022) provides access to the donors, samples, and datasets that have been collected and analyzed by HuBMAP teams. The EUI supports semantic and spatial exploration of the HRA, where all 55 organs and 1,203 tissue blocks can be viewed. However, identifying tissue blocks of interest is made difficult by occlusion in the small 2D interface, and by the challenge of navigating a 3D space on a 2D web browser.

#### (2) QA/QC by data providers

Registering tissue blocks into the HRA is an essential step for HRA construction and usage. However, errors in tissue registrations do occur, such as picking the wrong reference organ, e.g., left vs. right or male vs. female kidney. A quick check with specimen metadata associated with each tissue block can reveal existing data discrepancies, and a global visual check across all 5,000+ registered experimental datasets makes it possible to (a) quickly identify what spatial metadata is missing for how many organ-specific tissue blocks and (b) which tissue blocks are registered incorrectly, e.g., the size, location, or rotation of a tissue block is different from that a surgeon remembers. The HRA Organ Gallery makes it possible to quickly identify and correct wrongly registered tissue blocks.

### Apparatus

The HRA Organ Gallery v0.6 was developed using Unity 2021.3.11 (https://unity.com) for the open-source OpenXR platform (https://www.khronos.org/openxr). To facilitate interactions with the tissue blocks and organs, we use the XR Interaction Toolkit 2.0.4 (https://docs.unity3d.com/Packages/com.unity.xr.interaction.toolkit@2.0/manual/index.html). We targeted and optimized for the Meta Quest 2 with Oculus Touch controllers (https://www.meta.com/quest/products/quest-2). The majority of the development was performed on a Dell Precision 7560 running Windows 11 Pro, with 64 GB RAM, an 11th Gen Intel i9, 2.60 GHz processor, and a Nvidia RTX A2000 GPU.

We use the CCF API v1.0.0 (https://ccf-api.hubmapconsortium.org) to access the biological structure, spatial, and specimen data needed to construct the 3D scene from the CCF.OWL 2.1.0 (Herr et al., 2022).

## Results

### Current features

The HRA Organ Gallery v0.6 has a variety of features to support the semantic and spatial exploration of tissue blocks in the HRA (see **Figure 2**).

**Figure 2.**
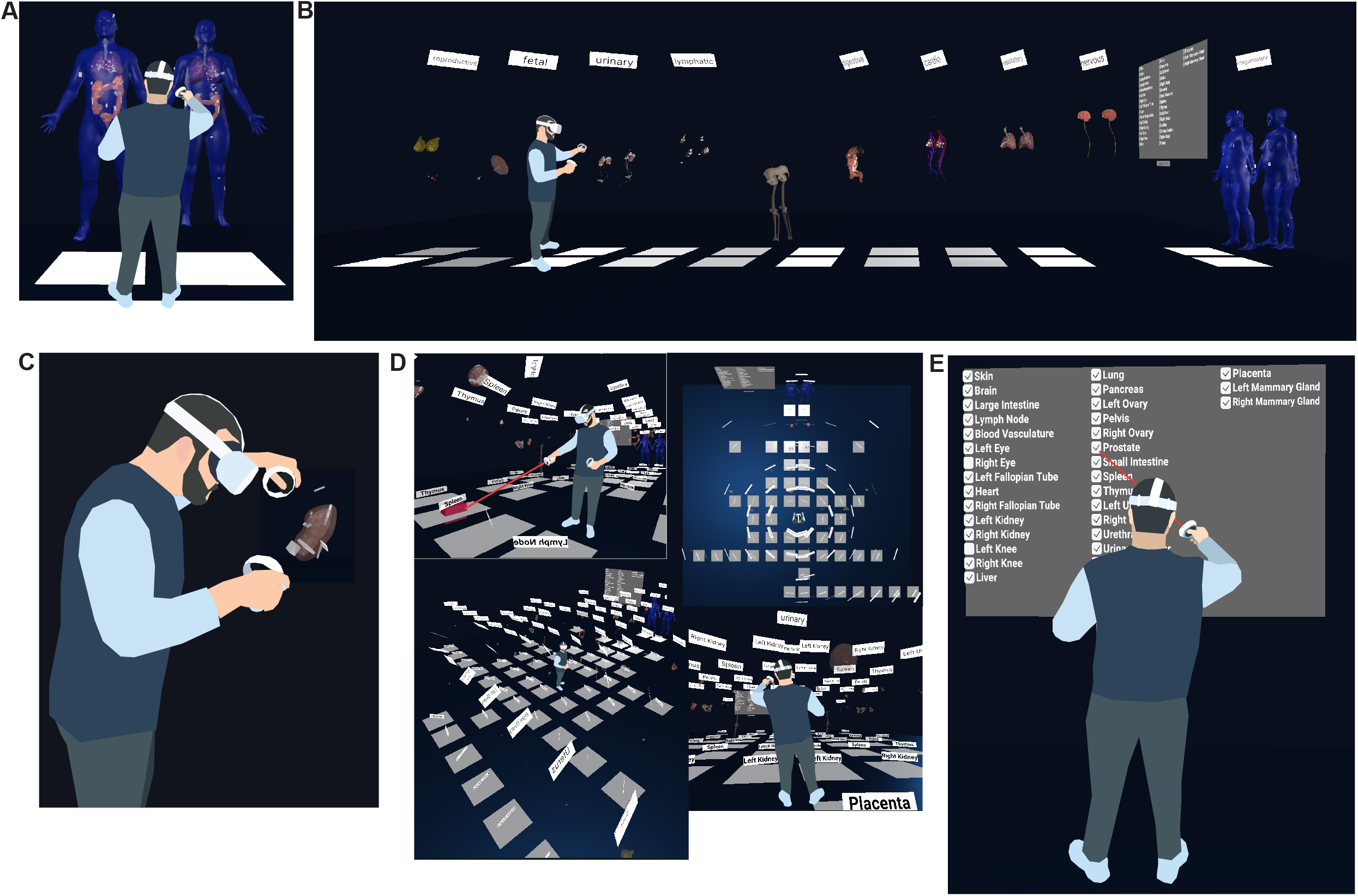
Functionality of the Organ Gallery v0.6. **(A)** Default organ layout **(B)** A user extrudes all 10 body systems containing 55 organs. **(C)** A user inspects a kidney and tissue blocks. **(D)** A user moves through the scene using teleportation, plus other views of the fully extruded gallery view. **(E)** A user employs the filter menu to show or hide organs.

#### Default organ layout (see **Figure 2A**)

All organs are arranged in the same fashion as in the EUI, where the male and female bodies appear next to each other, placed ~20 cm over the ground plane. All 55 organs in HRA v1.3 are in their anatomically correct locations inside the male and female skin. A filter menu, a console, and various UI elements provide functionality and information to the user. Labels for body systems and organs facilitate orientation and navigation in the scene.

#### Two-step extrusion system (see **Figure 2B**)

Once completely loaded, the user can pull out the organs along the global z-axis, first grouped by body system, and then individually. This results in a gallery-style grid. All registered tissue blocks stay with the organ. This makes it easy to spot wrongly registered tissue blocks when organs are moved, supporting the **QA/QC use case**.

#### Teleportation (see **Figure 2D**)

The user can perform natural movement in physical space, but to traverse the 30×30 ft floor (10×10 m), they can use teleportation using a red ray interactor. To minimize the time needed to arrive at the desired organ, white teleportation anchors on the virtual floor transport the user ~50 cm in front of the organ and adjust their orientation towards the organ when clicked.

#### Filter menu (see **Figure 2E**)

Showing all 55 organs at the same time creates performance issues for the targeted VR hardware. The default view in v0.6 thus only shows eight organs (skin, colon, lung, heart, all male and female). However, a filter menu on the left lets the user turn organs on and off through checkboxes.

#### Utilizing HuBMAP APIs (see **Figure 3**)

When loading, the app performs a call to the CCF API (https://ccf-api.hubmapconsortium.org/#/operations/scene). The JSON response is then deserialized into a custom C# class to capture relevant data fields, such as the tooltip name for organs and the list of anatomical structures with which tissue blocks collide. While 3D organ models are stored in local memory once the application has been run for the first time, it requires an internet connection to retrieve the most recent set of tissue blocks.

### Application architecture

**Figure 3** presents an overview of the most important building blocks of the application. When started, the */scene* endpoint of the CCF API is called, serving all available organs and tissue blocks as JSON objects. The centerpiece of the setup is the SceneBuilder (see **Figure 3B**), the data manager for the application. A series of C# classes retrieve data from the API and build the scene. The SceneBuilder references a SceneConfiguration with a URL to the CCF API. DataFetcher (see **Figure 3A**) defines a method to *Get()* this URL. To then receive the API response, DataFetcher uses three classes: Node, which contains fields corresponding to the body of the CCF API response; NodeArray, which wraps around the Nodes; and GLBObject, a container for the 3D organ to be loaded (in the GLB format). The SceneBuilder runs a suite of methods before or during the first frame or asynchronously: Methods in **orange** and **dark green** load and place organs and tissue blocks. Methods in **light green** add an OrganData component to each organ, which lets components across the app access its metadata. Methods in **pink** do the same for tissue blocks. Once completed, the user can start interacting with the organs. When running on the Meta Quest 2, it takes about 20 seconds for the entire scene to load.

**Figure 3.**
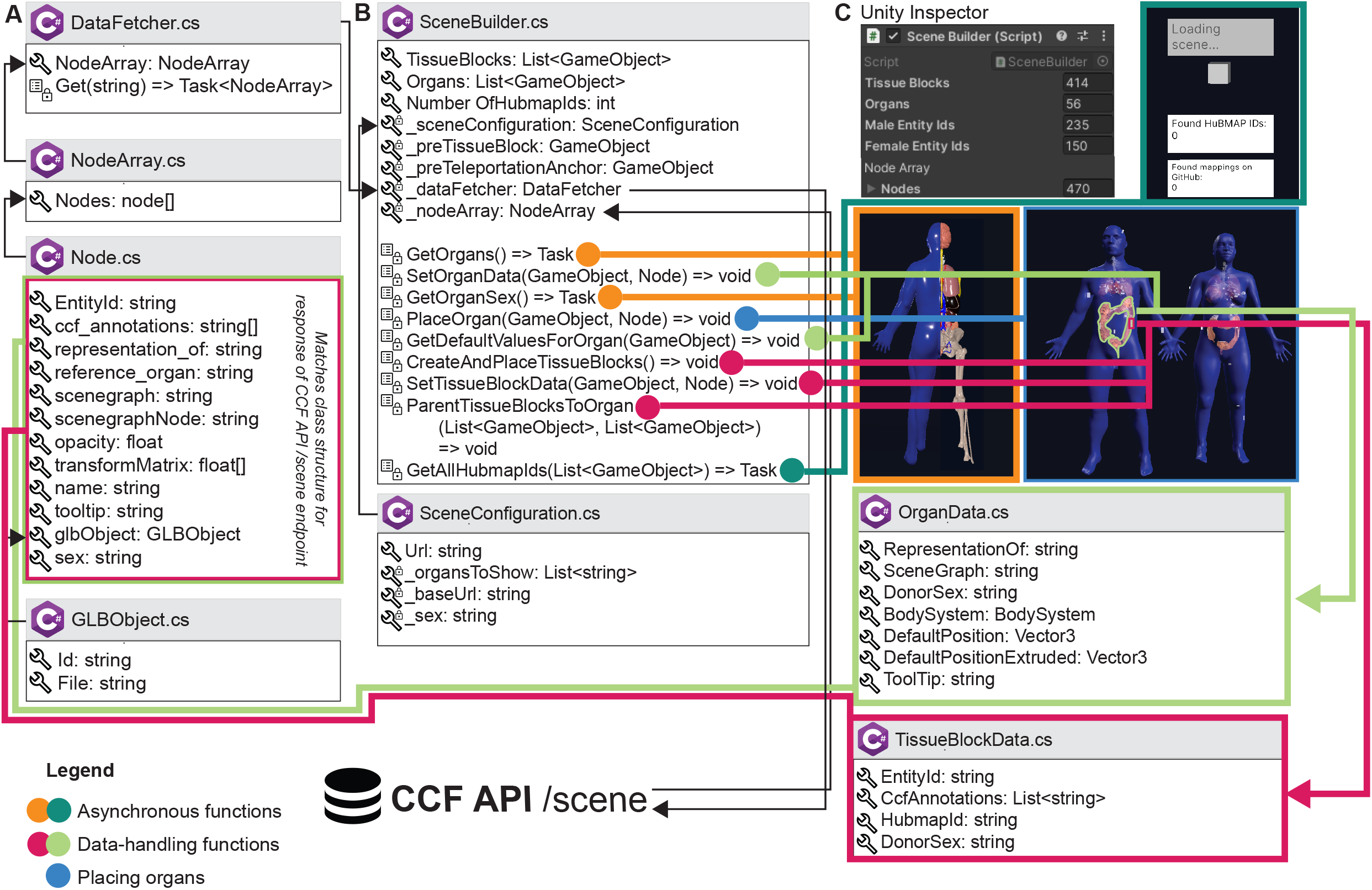
Architecture of the HRA Organ Gallery v0.6. Boxes with the C# logo signify a C# class or struct. A wrench symbolizes a property or field; a list symbolizes a method; private properties, fields, or methods have an added lock symbol. In some cases, we only show a selection of properties, fields, methods, and events for a class. *List<T>* and *Array[]* refer to collections and arrays. *Task* or *Task<T>* denotes an asynchronous function (with *<T>* meaning that a type is promised as return). **(A)** DataFetcher and its associated classes are used to retrieve data. **(B)** SceneBuilder sets up the 3D content. **(C)** SceneBuilder is derived from Unity’s native Monobehaviour class so it can be added to a GameObject in the Unity scene (**C, top left**). The Unity inspector then shows the SceneBuilder component, including displays the number of tissue blocks and organs received from the API, which is helpful for our developers. Note that the number of male and female entity IDs in **(C)** does not sum up to 414 tissue blocks, because the SceneBuilder captures sex only for HuBMAP tissue blocks.

### Planned features to support the two use cases

#### On-ramping

The goal of the UIs of the HRA is (1) to enable tissue providers to generate, verify, and (if needed) correct existing tissue block registrations, and (2) to allow HuBMAP Data Portal users to semantically and spatially explore datasets curated at high quality through unified pipelines. The HRA Organ Gallery aims to support this essential **on-ramping** use case with a series of VR superpowers.

##### On-Ramp-N1

As a <computational biologist or clinical researcher>, I want to <be able to pick tissue blocks with my hands in VR> so I can <examine the many anatomical structures that comprise them, explore what tissue blocks were registered where, and save relevant organs/blocks out for later examination>.

We are prototyping a **wrist pocket** (see **Figure 4A**), where the user can pick tissue block samples from an organ, deposit them into a virtual pocket tied to their VR controller, and then retrieve the metadata for the tissue block (such as its HuBMAP ID) via an API. After completing the VR session, the user will receive an email with links to all organs/blocks collected on the HuBMAP Portal. Lines between the tissue blocks on the wrist pocket and the reference organ help the user keep track of the original location of the tissue block.

**Figure 4.**
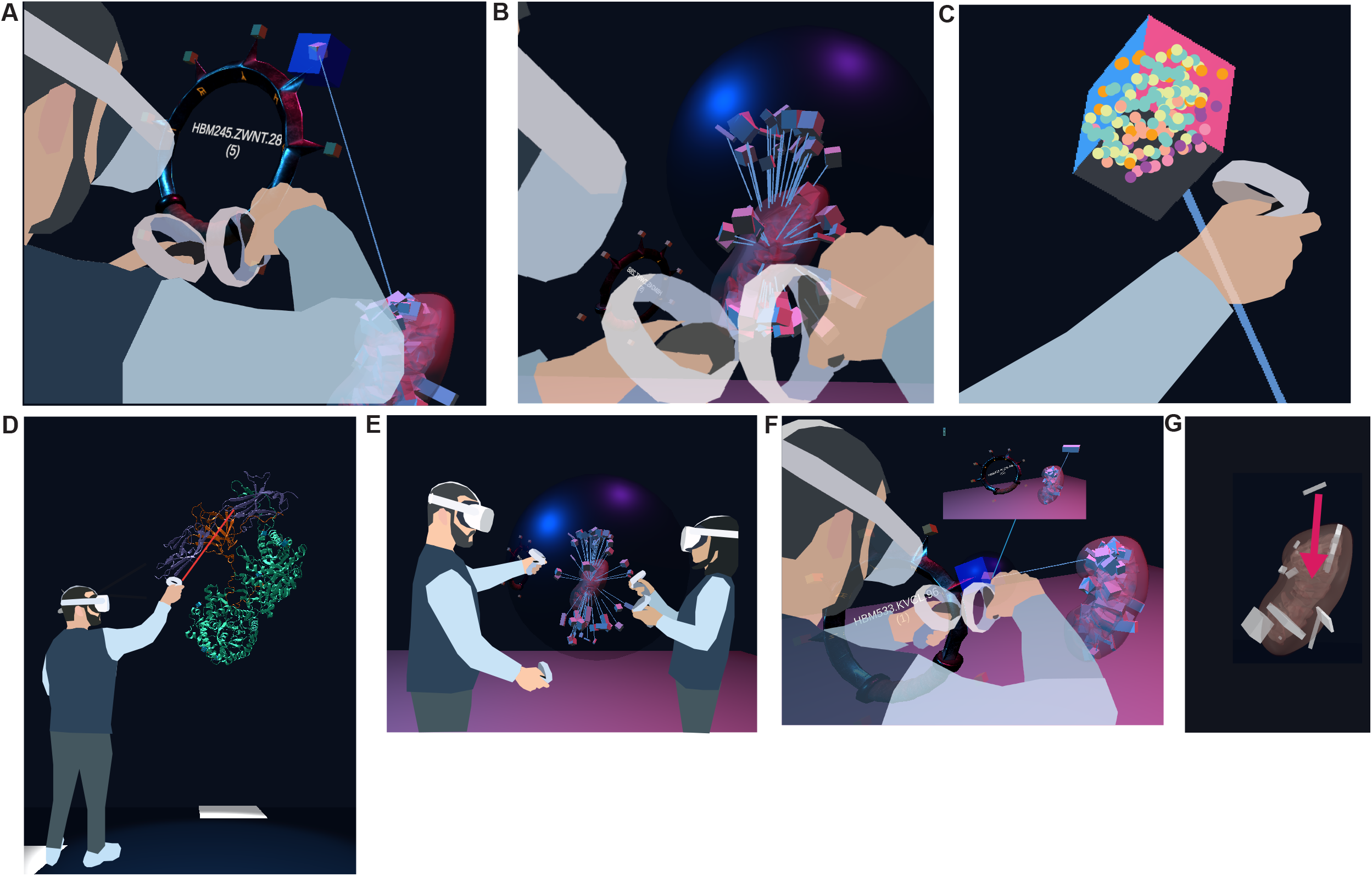
Features currently prototyped or planned. **(A)** A user picks a tissue block of interest and stores it in a wrist pocket. **(B)** A user disentangles overlapping tissue blocks with a probing sphere by extruding them from their shared centroid. **(C)** A user views the cell types inside a selected tissue block. **(D)** A user inspects an enlarged 5W21 protein. **(E)** Two users explore the HRA Organ Gallery together. **(F)** Bar graphs to visualize cell type counts for a selected tissue block. **(G)** A user corrects a misregistered tissue block by manually picking it up and moving it to the correct location.

##### On-Ramp-N2

As a <computational biologist or clinical researcher>, I want to <be able to examine multiple overlapping tissue blocks> so I can <understand the spatial distribution of tissue blocks in tight anatomical structures better>.

We developed a prototype of a **probing sphere** (see **Figure 4B**), tied to the user’s right hand, that can be used to hover over a set of overlapping tissue blocks to ‘explode’ the tissue blocks in a circular fashion away from the centroid of all the tissue blocks registered to the organ. We use collision detection between the sphere and the tissue block to enable the exploding motion for tissue blocks as needed. Lines connect the exploded position of the tissue block to its original position inside the organ. The size of the sphere and the degree of explosion can be adjusted by the user’s right and left thumb stick, respectively. The maximum distance between original and exploded position equals the maximum size of the sphere.

##### On-Ramp-N3

As a <computational biologist or clinical researcher>, I want to <move, rotate, and scale organs and the tissue blocks inside them> so I can <assess whether the reference organ fits the requirements for registering my tissue blocks>.

We are planning an **organ pullout feature**, where the user can grab, move, rotate, and rescale all 55 3D reference organs using the most natural input device: their hands (via VR controllers). This enables the user to switch between true, increased, and decreased scaling, default and custom rotation, and any position for the organ the user can reach with their hands. This enables the user to check if the reference organs will be a good fit for registering their tissue blocks. Additionally, it allows the user to inspect the organ and its tissue blocks from any angle–a non-trivial feature on the 2D interfaces of the EUI, where restrictions on camera movement prevent the user from overly experimental 3D camera movement. Note that all three functionalities described so far are neither available nor feasible in the EUI.

##### On-Ramp-N4

As a <computational biologist or clinical researcher>, I want to <see cell type populations for a selected tissue block> so I can <decide if a tissue block is relevant for my work>.

We are prototyping a **cell type population explorer** (see **Figure 4C**) that uses experimental data annotated via Azimuth (Hao et al., 2021) and ASCT+B tables to fill selected tissue blocks and anatomical structures with the correct number and type of cells in 3D. As we are using bulk data, the cell layout is random.

##### On-Ramp-N5

As a <computational biologist or clinical researcher>, I want to <view 3D models of proteins characterizing the cells of a selected tissue block> so I can <explore HRA data across scales>.

The planned **3D protein viewer** will allow experts to view selected proteins in VR (see **Figure 4D**). In the HuBMAP data ecosystem, protein biomarkers are recorded in the ASCT+B tables and in Organ Mapping Antibody Panels (OMAPs) (Radtke and Quardokus, 2021). The protein viewer will use data from NIH3D (Bioinformatics and Computational Biosciences Branch, 2023).

##### On-Ramp-N6

As a <newcomer to a HuBMAP team or a data provider>, I want to <be able to walk through the HRA in a co-op mode with another person> so I can <learn from a more experienced individual>.

The ability to collaborate in a **co-op mode** (see **Figure 4E**) will allow two or more users to walk through the scene, extrude organs, and inspect tissue blocks together. We will add annotation features and voice chat to facilitate communication between users.

### QA/QC

As of February 9, 2023, 19 users from 15 data providers have registered 1,203 tissue blocks from 292 donors using MC-IU’s forementioned RUI; out of these, 407 tissue blocks from 153 donors with 508 tissue sections and 1,421 datasets by 10 data providers have been published; all others are somewhere in the **QA/QC process**. While users register the spatial location of tissue blocks with great care, mistakes happen. Resulting errors in spatial or specimen data are difficult to identify when reviewing the data in 2D. However, they become immediately apparent in VR, where 3D organs and 3D tissue blocks can be examined in 3D up close from all angles and at different scales of magnification. Thus, the HRA Organ Gallery makes it possible for different end users (data curators, clinicians, researchers) to perform QA/QC using an intuitive and efficient VR user interface.

#### QA-QC-N1

As a <data provider>, I want to <see cell type populations for a selected tissue block> so I can <check if the tissue block was registered correctly>.

The 3D display of reference organs in VR can be enhanced by adding **auxiliary 2D data visualizations** of biological structure data (such as the number of cells per cell type) or specimen data (age, sex, BMI) for a tissue block of interest (see **Figure 4F**), based on previous work visualizing cell type distributions with stacked bar graphs (HuBMAP Consortium, 2022). We plan to build on the aforementioned concept of IRVEs (Bowman et al., 2003) to enhance the VR scene with ‘flat’ data visualizations. After initially focusing on tissue blocks and datasets from the kidney and skin, we aim to incorporate further ‘gold standard’ datasets as they become available. Utilizing the power of data visualization will enable users to quickly identify tissue blocks of interest, allowing them to make downstream or upstream adjustments in the HuBMAP data ecosystem based on ‘rogue’ tissue blocks with unusual cell type distributions.

#### QA-QC-N2

As a <data provider>, I want to <correct a wrongly registered tissue block> so I can <save time as opposed to correcting the registration in a separate 2D application>.

We are envisioning a **fix-on-the-spot feature** (see **Figure 4G**), where the user can perform a re-registration on the spot (e.g., from the left to the right lobe of the lung). This amended registration could then be assigned a permanent identifier, to be used in a variety of contexts (and even with other consortia). This feature would serve as a VR port of the RUI and combine two features of the existing CCF UIs (registration and exploration).

## Discussion

### Training materials

We will expand on the training and onboarding materials already created by producing a series of video tutorials and other documentation (written and in-app). These materials will guide and orient users, especially those unfamiliar with VR headsets, by explaining the controls and features in the application. They will also document how the HRA Organ Gallery ties into the evolving HRA and the data used to build it. We continue disseminating these training materials via YouTube and other social media platforms and will ship them as part of every release. We will explore deploying the HRA Organ Gallery to classrooms for anatomy and science students and the general public. This way, the HRA Organ Gallery becomes a valuable tool for HuBMAP outreach but also for ‘in-reach’ to train existing HuBMAP and other researchers working on the HRA. For example, we demonstrated the app at a Common Fund Data Ecosystem (CFDE) kickoff meeting in the Washington, DC area in November 2022, where we were able to show how tissue blocks from four different funded efforts (HuBMAP, the Genotype-Tissue Expression [GTEx] Project, the Kidney Precision Medicine Project [KPMP], and Stimulating Peripheral Activity to Relieve Conditions [SPARC]) can be harmonized, presented, explored, and queried within one coherent, immersive 3D space.

### Testing with experts

We will solicit further feedback from subject matter domain experts at the NIH, and its funded efforts. This will allow us to introduce and involve a wide variety of experts in the HRA effort, including experts in bioinformatics, 3D modeling, medical illustration, anatomy, data curation, and biology. We also plan a user study to gather quantitative data about performance for different QA/QC error detection methods and completion time and to test the usability, engagement, and presence afforded by the app.

## Conclusion

The HRA Organ Gallery provides an immersive view of HRA construction and usage, and with the organs and tissue blocks in their true size, enabling effective QA/QC using modern VR headsets and hand-held extended reality devices that are well documented and affordable (~$600 per set). It will soon be made available for free to more than 15 million VR devices (David Heaney, 2022)across the world via the Oculus Store (https://www.oculus.com/experiences/quest). We will develop extensive training materials (videos, written documentation, virtual walkthroughs inside the application) to enable a diverse array of users to utilize the HRA Organ Gallery for their own research and data curation.

## Acknowledgements

We would like to thank Kristen Browne and Heidi Schlehlein for creating the 55 3D models of the HRA v1.3. We deeply appreciate Bruce W. Herr II’s support in utilizing the CCF API. We are also grateful for Dena Procaccini’s contributions to existing and planned training materials. We thank Tabea Mader, Yongxin Kong, Naval Pandey, and Brent Iberg for appearing in our figures. Finally, we would like to acknowledge Gael McGill’s and Ushma Patel’s expert comments on the design goals and implementation of the HRA Organ Gallery. A preprint of this work exists on biorXiv.

We acknowledge funding from NIH awards OT2OD026671 (AB, KB) and OT2OD033756 (AB, SL, YRK, KB); Cellular Senescence Network (SenNet) Consortium Organization and Data Coordinating Center (CODCC) award #1U24CA268108-01 (AB, KB); Common Fund Data Ecosystem (CFDE) OT2 OD030545 (AB, KB); National Institute of Diabetes and Digestive and Kidney Diseases (NIDDK) Kidney Precision Medicine Project grant U2CDK114886 (KB); and Kidney Mapping Atlas Project (KMAP) grant 1U01DK133090-01 (KB).

## Contribution to the field statement

The work presented here is situated at the intersection of VR, information visualization, and bioinformatics. The HRA Organ Gallery enables users to explore 55 3D reference organs and 1,203 mapped tissue blocks from 292 demographically diverse donors and 15 providers with links to 5000+ datasets. The user can do so spatially and semantically in a coherent, immersive, 3D VR environment. First, prior intersecting work in these three research areas has focused on fewer organs (typically one or two) and has offered a less comprehensive look at data from donors in such numbers. Second, less value has been placed on advanced user interactions with the 3D organs and tissue block data in VR. Third, our work presents novelty in that user interfaces in most other cell atlassing efforts are mostly two-dimensional. Finally, to our knowledge, the outreach and in-reach potential of similar VR interfaces in other atlassing efforts has not been sufficiently explored.

## Data & code availability statement

The HRA Organ Gallery v0.6 was released on January 13, 2023, and is available to invited users via Oculus App Lab. We welcome feedback on the application and invite user testing, both in terms of engagement with this paper and with the VR app. All code is available at https://github.com/cns-iu/ccf-organ-vr-gallery under the CC-BY 4.0 license (https://creativecommons.org/licenses/by/4.0). The demo app is available for free via the Oculus Store (https://www.oculus.com/experiences/quest). Please reach out to lead author Andreas Bueckle if you are interested in being a test user. More information for test users is at https://github.com/cns-iu/ccf-organ-vr-gallery/blob/main/INFORMATION_FOR_TESTERS.MD. The CCF API can be accessed at https://ccf-api.hubmapconsortium.org.

## Author Contributions

AB and CQ are the corresponding authors. AB has led the writing effort for this article as well as the research and development for the HRA Organ Gallery. He also serves as the technical lead of the project. CQ wrote the related work sections and co-organized the article submission. SL contributed to the related work section. YRK served as co-developer of the HRA Organ Gallery until v.0.6. NP served as advisor for 3D models and application testing. KB supports the research and development effort for the HRA Organ Gallery as part of HuBMAP, she reviewed and edited the manuscript.

